# A Method for the Combined Functional Assessment of Photoreceptors and Ganglion Cells in Acute Retinal Explants

**DOI:** 10.1101/2024.12.03.626609

**Authors:** Xinyi He, Júlio César da Costa Moura, Pedro Gil Fernández, Paul Gläser, Lan Wang, François Paquet-Durand, Zhijian Zhao, Christina Schwarz

## Abstract

Significant progress has been made in studying the individual steps of retinal signaling, yet a comprehensive understanding of the functional interplay between photoreceptors, as input cells, and retinal ganglion cells (RGCs), as output cells, remains elusive. This connection is crucial to fully unravel the retina’s computational capabilities and its role in visual processing. To address this gap, this study introduces a novel technique that combines two-photon (2P) autofluorescence microscopy with multi-electrode array (MEA) recordings, enabling simultaneous testing of photoreceptor and RGC function. Unlike traditional methods that target isolated cell classes or structural analysis, this integrated technique provides the basis to investigate input-output relationships within the retinal neural network, offering a deeper understanding of its dynamic activity and adaptability to cellular dropout.

## Introduction

The retina is a sophisticated neuronal network responsible for initiating the processing of visual information. This process begins with phototransduction in rod and cone photoreceptors, where light is converted into electrical signals. Light activates the photopigments (e.g. rhodopsin and cone-opsins), prompting the transformation of 11-*cis* to all-*trans* retinal. This transformation triggers a biochemical cascade, including the activation of phosphodiesterase, which hydrolyzes cyclic guanosine monophosphate (cGMP). The reduction in cGMP levels leads to the closure of cyclic nucleotide-gated channels, resulting in photoreceptor hyperpolarization (Korenbrot, 2012). The electrical signals generated by photoreceptors are transmitted to bipolar cells and retinal ganglion cells (RGCs) for initial processing (Masland, 2012). RGCs are the output neurons of the retina that encode visual information through their spiking activity. With ∼40 subtypes, each type has a unique response to specific visual features (Baden et al., 2016; Bae et al., 2018; Rheaume et al., 2018; Goetz et al., 2022). This diversity enables the efficient relay of visual information to higher brain regions (Nguyen-Ba-Charvet et al., 2020; Johnson et al., 2021). Despite extensive research into the individual steps of retinal signaling, an integrated understanding of the functional interplay between retinal input (photoreceptors) and output (RGCs) remains elusive. Investigating this connection is crucial for uncovering the retina’s full computational capacity and its role in vision.

One example where correlated functional measurements of input and output retinal cells are of interest, are studies of visual processing in degenerating retina. For instance, in the context of retinal degeneration, a recent study suggests an unexpected resilience of the retinal network in rd10 mice. Despite the loss of approximately 90% of photoreceptors, RGCs appear to maintain functional activity (Dyszkant et al., 2024), suggesting a remarkable adaptability of the system.

Current methods to study and evaluate retinal function include immunohistochemistry, electrophysiology, and imaging, employed either individually or sequentially (Mead et al., 2016; Hasegawa et al., 2016; Molday et al., 2019; Rubin et al., 2022; Leinonen et al., 2022; Tebbe et al., 2024). While electrophysiological techniques such as patch-clamp recordings and multielectrode arrays (MEAs), (Meister et al., 1994; Charvet et al., 2010; Shabani et al.,2021; Goetz et al,. 2022) as well as two-photon (2P) imaging of sensors, such as calcium or voltage indicators (Borghuis et al., 2011; Baden et al. 2016; Xu et al.,2023; Baez et al., 2024), are highly effective in assessing RGC function, they often fail to capture the functional input from photoreceptors. In such cases, photoreceptor function is typically inferred from morphological data obtained through immunostaining for cell numbers and functional proteins (Roche et al., 2016; Yan et al., 2024) or is measured separately through imaging of photoreceptor axon terminals using biosensors (Wei et al., 2012; Chapot et al 2017; Szatko et al 2020).

To address this methodological gap, we propose a novel approach to correlate photoreceptor and RGC function in acute retinal explants using a combination of MEA and two-photon autofluorescence recordings. Our method is suitable to study signal integration and neural encoding within the retina, providing a tool to investigate both normal retinal function and the impacts of cellular degeneration, which could provide guidance for future therapeutic interventions.

## Methods

### Combined setup for 2P microscopy and MEA recordings

A customized 2P microscope was modified by adding an imaging platform for heavy loads (Figure 1), enabling recordings from RGCs using a MEA and from photoreceptors with 2P autofluorescence microscopy. In brief, the microscope is driven by a Ti:Sapphire laser (MaiTai XF-1 DS, Newport Spectra-Physics) and equipped with a 16x water immersion objective (CFI75 LWD 16X W, Nikon,) and two detection channels for simultaneous data acquisition. In this study, the pulsed laser was set to 730 nm emission to target endogenous functional fluorophores (Figure 1C). One channel recorded reflected light, and the other channel recorded two-photon excited autofluorescence (400-570 nm, beamsplitter: ZT485DCRB + Short-pass filter: 633SP, AHF). We used the commercially available electronic and software solution FidelasHexSetup (Brightview Robotics). In order to fit the MEA head stage with the imaging system, an XY-movable table (8MTF-102LS05, Standa) and a heavy-load Z-translation stage (MLJ150/M, Thorlabs) are installed below the objective to position the sample. An additional piezo motor is mounted together with the objective holder, for fine tuning in Z-direction. The motorized stage system is controlled through a python script and operated via a game controller (516L8AA, HyperX). The script is available on GitHub via the link below: https://github.com/AuSchwarzLab/2P-imaging-and-MEA.

**Figure 1.**
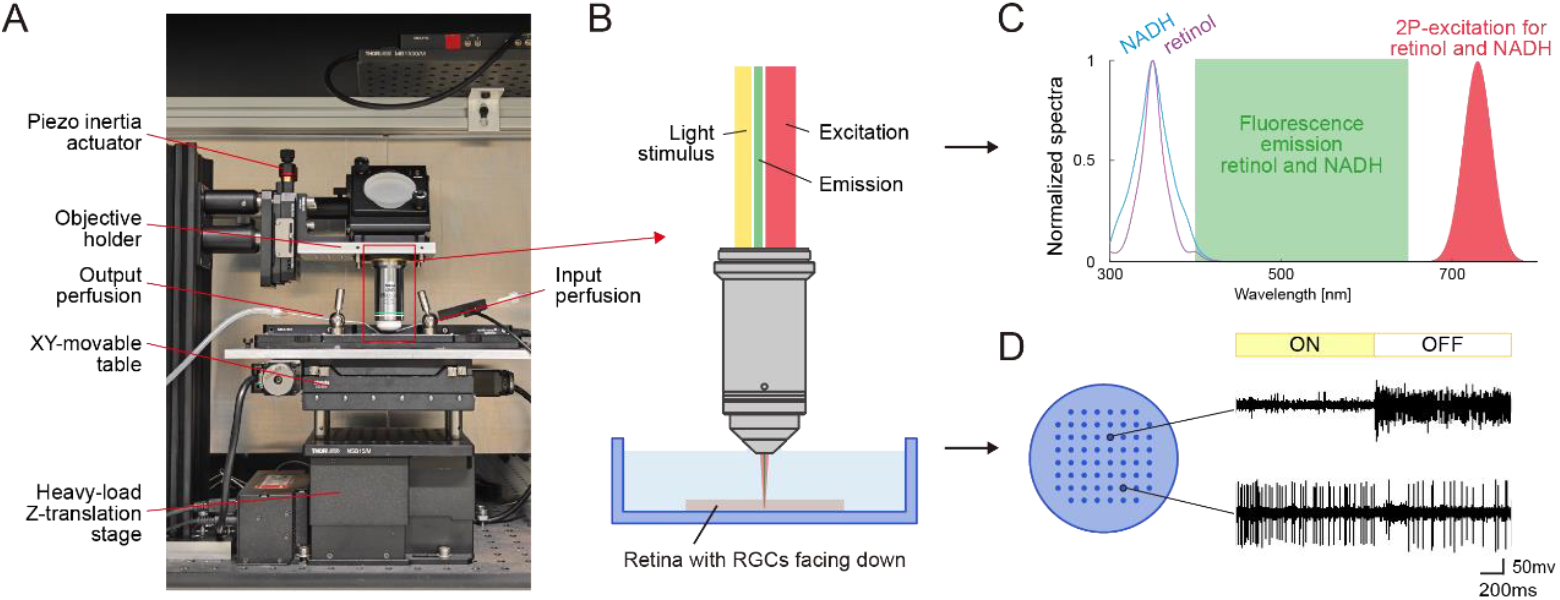
Overview of the two-photon (2P) microscope together with the multi-electrode array (MEA) device. (**A**) Photograph of the head stage combining the 2P-microscope and the MEA device. (**B**) Illustration of the 2P-imaging and MEA-recording configuration. (**C**) Normalized excitation spectra of endogenous fluorophores (all-*trans*-retinol and NADH) within photoreceptors, the illuminating wavelength of the pulsed laser and the spectral band that is captured by the fluorescence channel. (**D**) Illustration of the MEA recording to capture the electrical activity of RGCs. Exemplary traces indicate the activity of two distinct RGCs upon ON-OFF light stimulation.

### Animals and tissue preparation

All animal experiments were approved by the institutional animal welfare committee of the University of Tübingen and performed according to guidelines (protocol AK04/22M). The study utilized retinas from adult C57Bl/6J mice (wild-type, JAX 000664, N = 3/6 mice/retinas) of either sex.

Animals were housed in the local animal facility under the standard 12h/12h day/night cycle at 22°C with 55% humidity. Before the experiment, animals were dark-adapted for >1 hour, followed by anesthesia using isoflurane and sacrificed through cervical dislocation. The eyeballs were carefully removed by cutting the optic nerve and surrounding tissues. Enucleated eyes were dissected in carboxygenated (95% O_2_ / 5% CO2) AMEs medium (Sigma, pH 7.4). The anterior parts of the eyes were removed, and the retinas were carefully isolated from the eyecups. For the experiment, the retinas were transferred to an MEA recording chamber so that the RGC layer faced the electrodes (Figure 1B). The recording chamber was perfused with carboxygenated AMEs medium at 37°C. All procedures were carried out under very dim red (>650 nm) light.

### Electrophysiology on ganglion cells

In this study, we used MEAs with a 180 µm thin recording area, containing 59 recording electrodes in an 8 × 8 configuration (60MEA200/30iR-Ti-pr-T type MEAs and MEA2100 system, Multi Channel Systems). The electrodes measured 30 µm in diameter and were spaced 200 µm apart. Before the measurement, the tissue was allowed to adjust to the experimental environment for ∼20 min so that neuronal activity stabilized. Spontaneous RGC activity was recorded for 30 s followed by 30 s of light-evoked activity during well-controlled light stimuli (see details below). The MEA system was operated using MC-Rack software (Multi Channel Systems), configured with a sampling rate of 50 kHz.

All electrophysiology data were spike-sorted using SpyKING Circus (Yger et al., 2018). Comprehensive details about the algorithm and its implementation are available on their webpage (http://spyking-circus.readthedocs.org/). The resulting spike clusters were inspected one-by-one and refined using the graphical interface of the phy package (http://phy.readthedocs.org/). During this process, clusters were manually merged or split based on the analysis of autocorrelograms, cross-correlograms and feature drift across the recording duration. As the output of spike sorting, responses of individual RGCs were assigned to each cluster. To assess how effectively a cell responded to a given stimulus, we computed the signal-to-noise ratio as described previously (Baden et al. 2016):

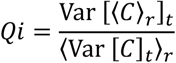

where *C* is the *T* by *R* response matrix (time samples by stimulus repetitions) and ⟨ ⟩_*x*_ and *Var*[]_*x*_ denote the mean and variance across the indicated dimension, respectively. For further analysis, we only used cells that responded well (Qi>0.5).

From the firing traces, RGCs were classified ON-cells, ON-OFF-cells, and OFF-cells based on the contrast in stimulus-evoked response (Baden et al., 2016; Carcieri et al., 2003):

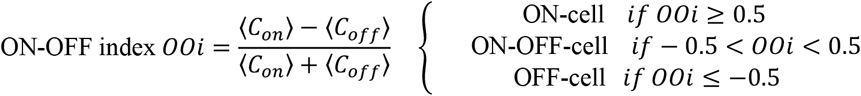

where ⟨*C*_*on*_⟩ and ⟨*C*_*off*_⟩ are all responses during the “light on” and “light off” phases, respectively.

### Light stimulation

A green LED (590 nm, Thorlabs) was selected for stimulation of both M-cones and rods of the mouse retina during electrophysiology. Full-field flicker stimuli (1 mm in diameter) at 0.5 Hz were delivered at an intensity of 1.01×10^12^ photons/s (R_Iso, M-cone_ = 1733.03 P^*^cone/s, R_Iso,rod_ = 4332.57 P*/rod/s) through the objective to the photoreceptors (Figure 1B).

For functional imaging of photoreceptors, the retina was either dark adapted for 10 min or bleached with the green LED set at 1.34 µW for 3 s. Notably, the imaging laser also stimulates the photoreceptors due to a 2P effect (Euler et al., 2009; Palczewska et al., 2014; Sharma et al., 2016) during the recording.

### Imaging of photoreceptors

Immediately following MEA recording, the tissue was imaged at the photoreceptor segments. Reflectance images provide information as to the morphology of the cell compartments, whereas two-photon excited autofluorescence refers to functional activity (Sharma et al., 2016; Sharma et al., 2016; Sharma et al., 2017, Walters et al., 2018). A functional photoreceptor should support the visual cycle in its segments to restore their sensory function after bleaching. All-*trans*-retinol, one of the intermediate products of visual cycle, is strongly fluorescent, providing an endogenous probe for testing the functionality with imaging techniques (Huynh et al., ARVO 2022, Nguyen et al., 2024). Fluorescence videos were captured for 60 s at a rate of 26.7 Hz. The imaging field was 80 µm × 80 µm. Both laser power and PMT gain were maintained when the tissue was imaged during dark adaptation and following a strong bleach. Photoreceptor videos were analyzed using ImageJ software (NIH).

## Results

As a first step RGC function was tested. The light stimuli triggered responses in ∼ 26 RGCs per section tested. Out of these cells, 25 RGCs presented stable responses. In the total population of 25 cells, we recorded 11 ON-, 9 ON-OFF-, and 2 OFF-cells. The remaining cells did not exhibit a typical light response pattern but were identified as nonclassified RGCs. RGC responses of four exemplary cells, i.e. two ON-cells, two ON-OFF cells and one OFF-cell, are shown in Figure 2. Although the ON-OFF index in Figure 2L is below 0.5, this cell is still categorized as an ON transient cell based on its light response patterns.

**Figure 2.**
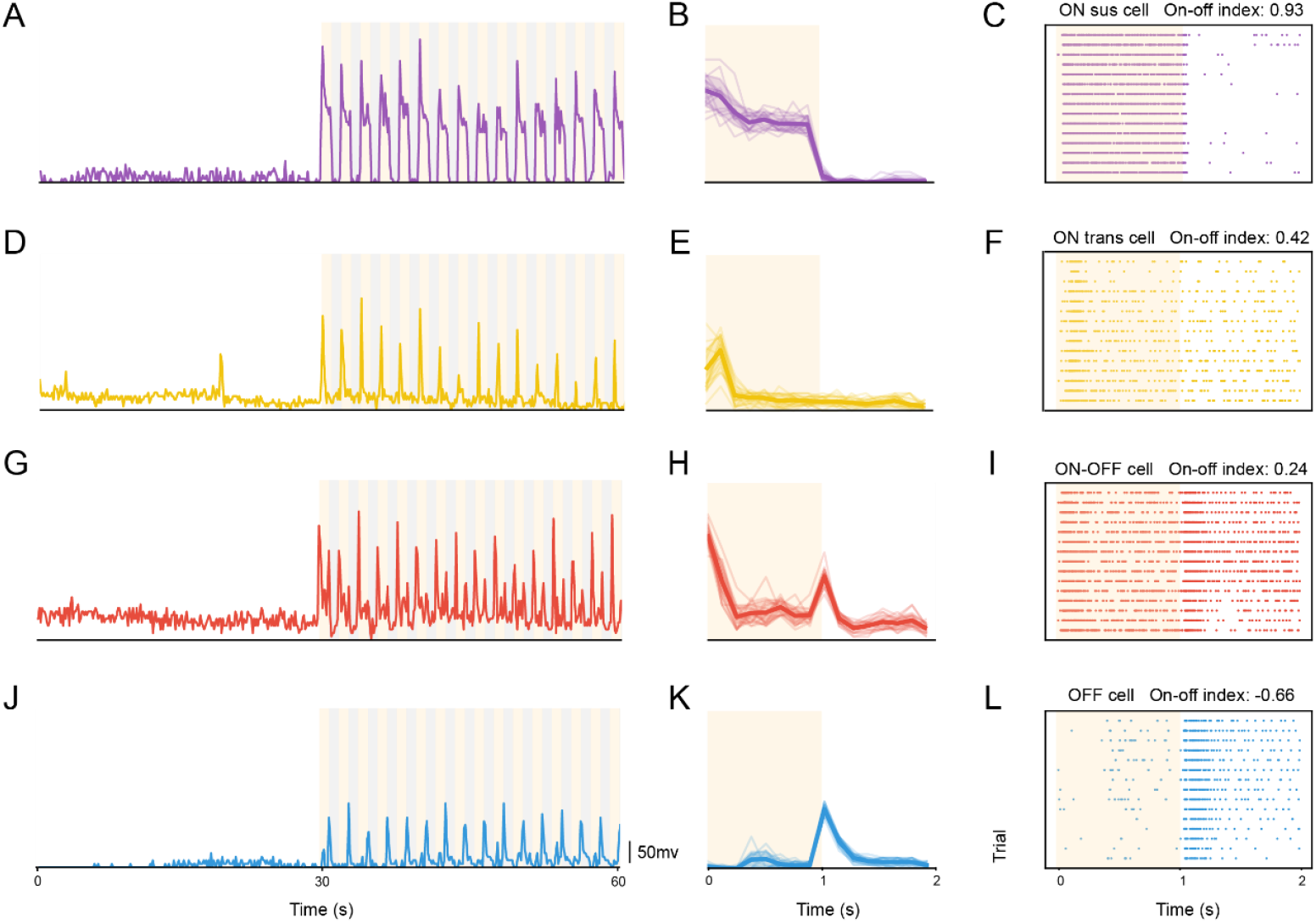
MEA recordings of RGC function. **(A, D, G & J)** Firing traces of four exemplary RGCs under light stimulation. (**B, E, H & K)** average firing patterns across all light stimulus snippets. (**C, F, I & L)** raster plots with corresponding ON-OFF index, supporting the functional RGC types.

In addition to functional properties of individual RGCs, the MEA data also revealed the electrode that recorded these spike trains. By imaging individual electrodes and recording the photoelectric artifacts, each electrode’s position could be translated to the coordinates of the motorized XYZ-stage.

Approximately 180 µm above the MEA electrodes, largely intact photoreceptor mosaics could be captured at the segment planes (Figure 3A). The photoreceptor segments are displayed in *en face* view at sequential focal planes 3 µm apart, stepping from outer towards inner segments. Individual functional photoreceptor segments are distinguishable based on their autofluorescence profile. At the highest focal plane (z= 0 µm), the mosaic is composed of cells of equal size, approximately 1.30 µm in diameter, and is interrupted at several locations by gaps. As the focal plane shifted towards the inner retina (from left to right), some gaps gradually revealed cells that were larger in diameter (∼2.76 µm), likely to be cone segments. The average functional photoreceptor density was approximately 2.9×10^5^/mm^2^ across the 6 retinas.

**Figure 3.**
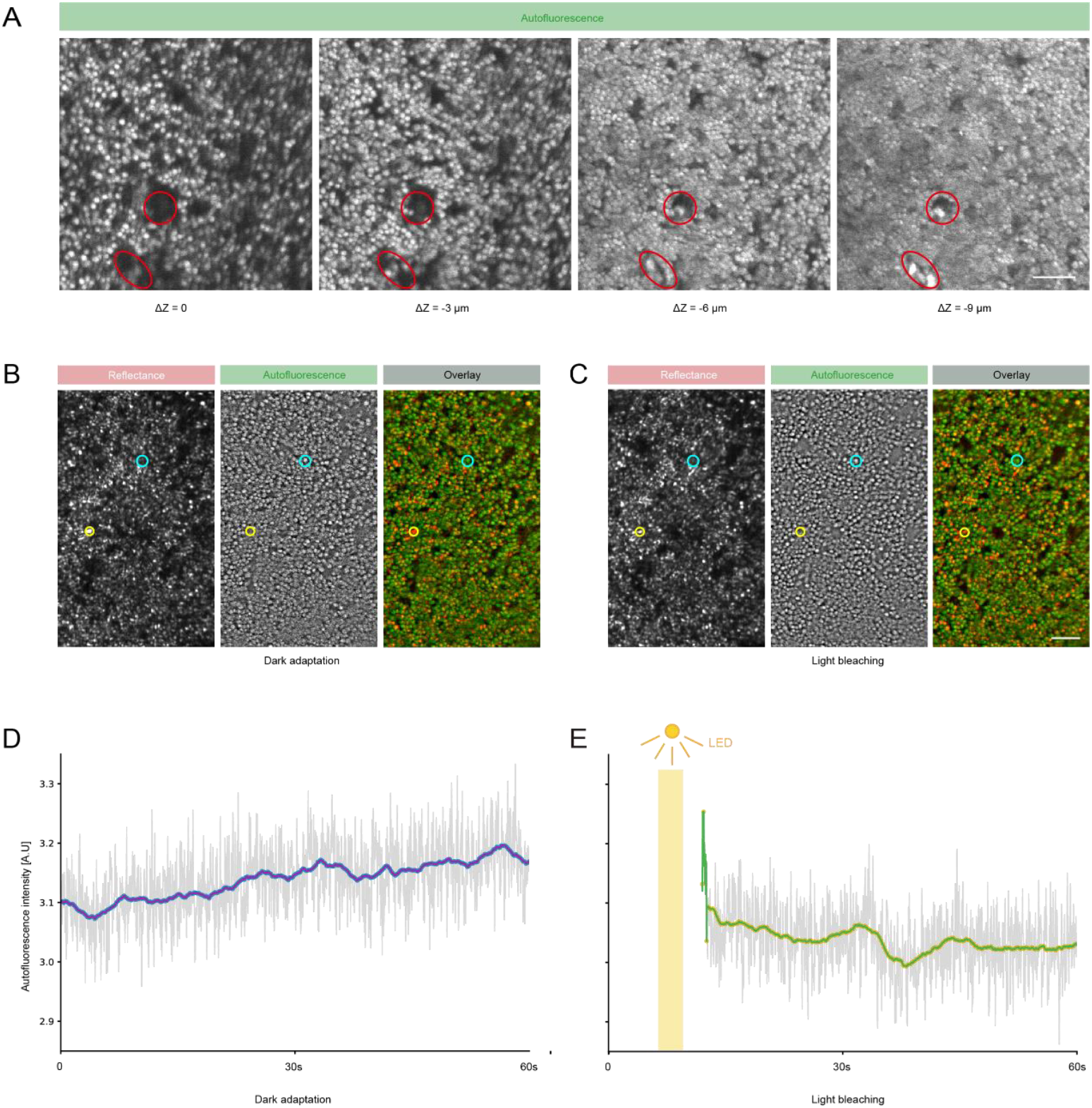
Functional 2P-imaging of photoreceptors. (A) Photoreceptor segments were revealed by 2P-imaging for autofluorescence. (B & C) Functional imaging for photoreceptors following 10 min of dark adaptation and 3 s of light bleaching with reflectance images (left), 2P autofluorescence images (center) and overlays (right). A subset of cells shows strong reflectance, but weak fluorescence (yellow circle), indicating morphologically intact but non-functional outer segments. Another subset of cells is indistinguishable in reflectance, but emits fluorescence (cyan circle), suggesting non-waveguiding yet functional outer segments. (D & E) Average time course of fluorescence following dark adaptation and light bleaching, respectively. (Scale bar: 10 µm)

Finally, we tested the retina for photoreceptor function after dark adaptation (Figure 3B) and following intense, instantaneous bleaches (Figure 3C). In corresponding reflectance and autofluorescence images, most photoreceptor cells exhibited a clear one-to-one correlation between the two channels. However, approximately 4.24% of the photoreceptors (e.g. yellow circle) were identifiable in reflectance but lacked substantial autofluorescence, suggesting that these cells were morphologically intact but lacked functional activity. Conversely, ∼10.65% of the photoreceptors (e.g. cyan circle) displayed pronounced autofluorescence but were not visible in reflectance. This pattern implies that these cells were functionally active and likely morphologically intact, but with suboptimal reflectance properties.

After dark adaptation, the frame-averaged autofluorescence intensity slowly increased when the recording started (Figure 3D), reflecting the stimulation by the imaging laser. Following an intense bleach (Figure 3E), autofluorescence intensity was high initially and quickly declined to a base line level. The transient elevation immediately after light bleaching is likely attributable to the increased levels of all-*trans*-retinol generated during light stimulation.

## Discussion

Retinal function and information processing have been extensively studied through electrophysiology and imaging. Most of the studies were limited to one particular cell type to understand its role or function within the network. Here, we introduced a combined method of functional 2P-microscopy and MEA recordings to correlate photoreceptor and RGC function. With further refinement, this approach could provide data to establish the input-output relationship of the retinal network under both normal and degenerative conditions.

We initiated our experiments by testing the function of RGCs. Based on the cells’ light responses, we could not only determine the functional properties of different RGCs, but also estimate their approximate locations. By correlating the photoelectric artifacts visible on the MEA with reflectance images, we accurately aligned the imaging area with specific MEA electrodes. While we employed a simple full-field flicker stimulus, more sophisticated paradigms, such as multi-color chirp and patterned stimuli, could enable comprehensive functional classification of RGCs. These approaches would also allow for precise mapping of their receptive fields, revealing both the number and specific types of photoreceptors that contribute input to each RGC.

Importantly, photoreceptor imaging can also serve as a quality control of sample preparation. A frequent critique of retinal explants is that outer segments may be lost during preparation. We calculated the photoreceptor density in the fluorescent image, which is approximately 2.9×10^5^/mm^2^ in the segment layer. This number aligns with those of a previous study (rod: 4.37×10^5^ cells/mm^2^, Cone:1.24×10^4^ cells/mm^2^) (Jeon et al., 1998). The difference in segment length and diameter also agrees with previous data (Ozaki et al., 2020). Distinguishing between rods and cones in *en face* images of mouse photoreceptors poses a significant challenge, as their morphological differences are subtle and not easily discernible, especially in reflectance images (Figure 3B & 3C).

A further advantage of our imaging approach lies in its ability to utilize autofluorescence to assess photoreceptor function directly. While similar data can be obtained using a combination of MEA recordings and reflectance imaging of photoreceptors, the latter is inherently less informative. Photoreceptor reflectance is strongly dependent on their waveguiding ability. In our study, some photoreceptors – though autofluorecent – did not produce detectable reflections, likely due to bending.

Imaging endogenous fluorophores associated with photoreceptor function has been previously demonstrated in living primates (Hunter et al., 2010; Sharma et al., 2016; Sharma et al., 2017) and recently in rabbits (Nguyen et al., 2024). Here, for the first time, we show similar outcomes in mouse photoreceptors of acute retina explants. However, our experimental setup differs from *in vivo* conditions in several critical ways. For instance, the absence of blood flow, which could reduce oxygen availability, is mitigated by AMES perfusion. Additionally, the removal of the retinal pigment epithelium (RPE) disrupts the traditional visual cycle, which further influences photoreceptor function by slowing the visual cycle. Despite these differences, our autofluorescence-based approach proves that all-*trans*-retinol can be produced and removed from the photoreceptor segments and provides thus a robust and meaningful assessment of photoreceptor functionality.

Alternative approaches to ours may include simultaneous imaging of two retinal layers using biosensors. However, this approach may alter the light sensitivity of photoreceptors due to the high-intensity light required for imaging, potentially affecting the accuracy of the recorded responses. Another commonly used method is electroretinography (ERG), which measures the summed electrical responses of retinal layers to light stimulation. While ERG provides insights into overall retinal function, it lacks the resolution to study activity at the level of individual cells, making it unsuitable for detailed input-output analyses. We demonstrated that our method has significant potential to address this gap. A logical next step would be to investigate how photoreceptor loss impacts the remaining retinal network during degeneration. By analyzing the input-output relationship, this approach could uncover the mechanisms underlying the retina’s remarkable ability to maintain functional robustness despite cellular degeneration.

## Notes

### Competing Interest Statement

The authors have declared no competing interest.

